# hogwash: Three Methods for Genome-Wide Association Studies in Bacteria

**DOI:** 10.1101/2020.04.19.048421

**Authors:** Katie Saund, Evan S. Snitkin

## Abstract

Bacterial genome-wide association studies (bGWAS) capture associations between genomic variation and phenotypic variation. Convergence based bGWAS methods identify genomic mutations that occur independently multiple times on the phylogenetic tree in the presence of phenotypic variation more often than is expected by chance. This work introduces hogwash, an open source R package that implements three algorithms for convergence based bGWAS. Hogwash additionally contains two burden testing approaches to perform gene- or pathway-analysis to improve power and increase convergence detection for related but weakly penetrant genotypes. To identify optimal use cases, we applied hogwash to data simulated with a variety of phylogenetic signals and convergence distributions. These simulated data are publicly available and contain the relevant metadata regarding convergence and phylogenetic signal for each phenotype and genotype. Hogwash is available for download from GitHub.

**DATA SUMMARY:**

1. hogwash is available from GitHub under the MIT license (https://github.com/katiesaund/hogwash) and can be installed using the R commands

~~~
install.packages(“devtools”)
devtools::install_github(“katiesaund/hogwash”)
~~~
2. The simulated data used in this manuscript and the code to generate it are available from GitHub (https://github.com/katiesaund/simulate_data_for_convergence_based_bGWAS)

**IMPACT STATEMENT:** We introduce hogwash, an R package with three methods for bacterial genome-wide association studies. There are two methods for handling binary phenotypes, including an implementation of PhyC(1), as well as one method for handling continuous phenotypes. We formulate novel indices quantifying the relationship between phenotype convergence and genotype convergence on a phylogenetic tree. These indices shape an intuitive understanding for the ability of hogwash to detect significant intersections of phenotype convergence and genotype convergence and how to interpret hogwash outputs.

## INTRODUCTION

### Bacterial Genome-Wide Association Studies

Bacterial genome-wide association studies (bGWAS) infer statistical associations between genotypes and phenotypes. Seminal bGWAS papers identified novel variants associated with antibiotic resistance in *M. tuberculosis* and host specificity in *Campylobacter*(1,2). Since then, there have been numerous applications of bGWAS that have further highlighted the potential of this approach to identify genetic pathways underlying phenotypic variation and provide insights into the evolution of phenotypes of interest. Association studies can use various genetic data types including single nucleotide polymorphisms (SNPs), k-mers, copy number variants, accessory genes, insertions, and deletions. To improve the power and interpretability of bGWAS inclusion criteria or weighting can be applied to these variants based on predicted functional impact, membership in pathways of interest, or other user preferences(3,4). Differences between human and bacterial GWAS have been reviewed extensively by Power *et al.*(5). Of note, clonality and horizontal gene transfer complicate the application of human GWAS methodology to bacteria. However, bGWAS approaches can leverage unique features of bacterial evolution, including frequent phenotypic convergence and genotypic convergence, to identify phenotype-genotype correlations.

### bGWAS Software

Several different variations of bGWAS approaches have been applied, including methods for SNPs, accessory genes (Scory)(6), or k-mers (pyseer) (7), methods using regression (pyseer)(7,8) or phylogenetic convergence (PhyC, treeWAS)(1,9), and methods designed for humans (PLINK)(10) or specifically for bacteria(7,9). Differences between regression based and convergence based bGWAS were expertly reviewed by Chen and Shapiro(11). Convergence based methods identify events where a genomic mutation arises independently on different edges of a phylogeny more often in the presence of the phenotype of interest than expected by chance (Figure 1C). Convergence based methods can yield higher significance with a smaller sample size, but may fail to identify some statistical associations that traditional GWAS approaches would identify when the population is clonal(11). Additionally, convergence based methods are limited to smaller data sets because of their large memory requirements and computational time relative to traditional methods(12), but can surmount issues of clonality.

**Figure 1.**
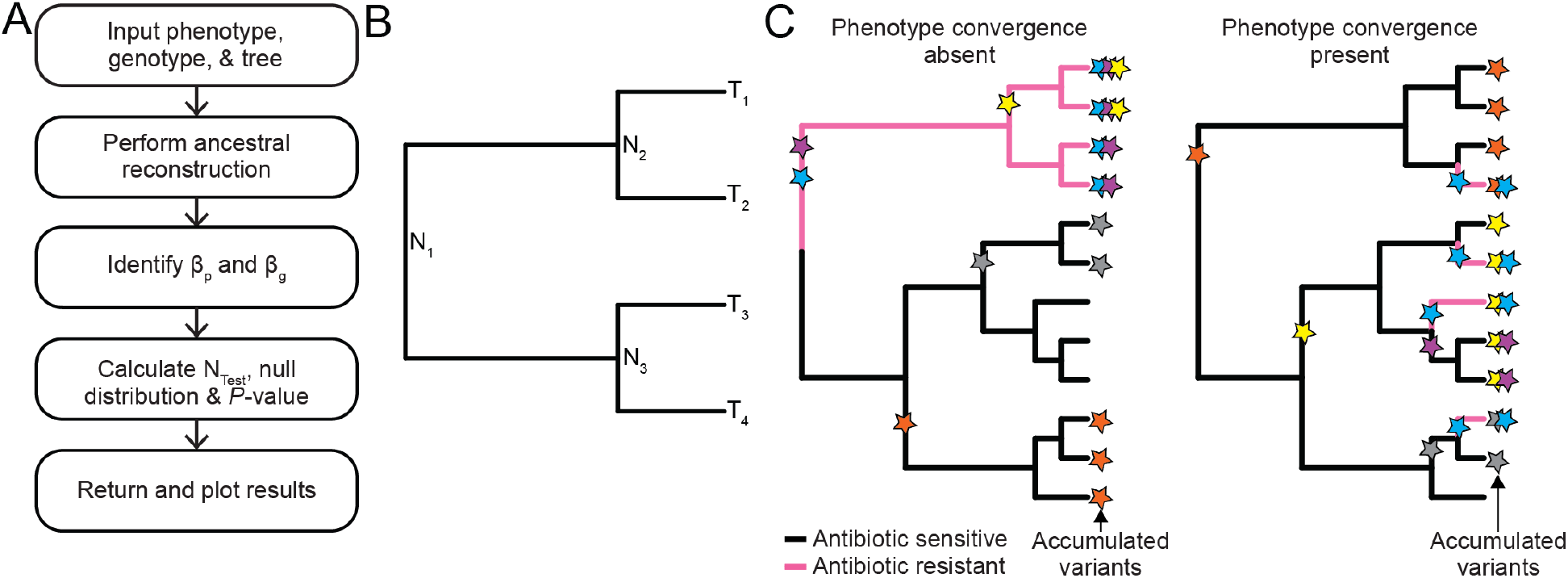
Hogwash workflow, tree nomenclature, and convergence example. A) Software workflow. B) In this example phylogenetic tree N_1_ is the root. Tree nodes are labeled N_1_ – N_3_. Tree tips are labeled T_1_ – T_4_. N_1_ is a parent node to N_2_ and N_3_. N_2_ is a child of N_1_ and a parent to T_1_ and T_2_. Edges are lines connecting a parent node to a child node or a parent node to a tip. C) A conceptual example of a phylogenetic tree with a phenotype that has arisen under two different scenarios. In the left tree antibiotic resistance, encoded by pink edges, arises once and therefore does not converge on the tree. In the right tree antibiotic resistance arises four times and therefore converges. Each colored star represents a unique genomic variant, such as single nucleotide polymorphism (SNP), that arises. Stars on edges indicate the time at which the SNP is inferred to have arisen. The stars at each tip indicate the accumulated variants found in each sample. In both trees the blue variant occurs in 4/4 antibiotic resistant isolates and 0/8 antibiotic sensitive isolates. Convergence based association methods could only ascertain the relationship between the blue variant and antibiotic resistance in the right tree.

### Objective

As the popularity of bGWAS increases there is a need to have more widely available software that addresses specific aspects of bacterial evolution and is appropriate for various kinds of datasets. This work introduces two novel methods for convergence based bGWAS with these needs in mind: the Synchronous Test and the Continuous Test. Users can implement these methods using hogwash, a new R package available on GitHub. Hogwash also contains an implementation of PhyC which is a bGWAS algorithm introduced by Farhat *et al.*(1). The Synchronous Test is a stringent variation of PhyC, requiring a tighter relationship between the genotype and phenotype. We describe the algorithms and evaluate them on a set of simulated data. The hogwash wiki contains further explanation of bGWAS, a more conceptual introduction to these three algorithms and specific user instructions for hogwash on a set of data provided with the software package (https://github.com/katiesaund/hogwash/wiki).

### Grouped Genotype Analysis

Some phenotypes are not well correlated with commonly occurring genomic variants. In these cases, rare variants may provide some additional explanation for trait variability. There are multiple approaches to studying rare variants including various burden testing methods which can group loci into meaningful groups, such as mapping SNPs to genes(13,14). Analyzing aggregated loci can improve both the interpretability of GWAS results and improve power to detect associations(13–16). Hogwash implements two such grouping approaches to improve convergence detection for related but weakly penetrant genotypes.

### Data Simulation

We evaluate hogwash results on simulated data generated to capture aspects of bacterial evolution pertinent to these bGWAS approaches. We simulated data with a range of phylogenetic signals and convergence distributions to highlight the critical impact of these features on bGWAS results. The simulated data are publicly available and could be used to compare the impact of convergence patterns within phenotypes, genotypes, and their intersection when benchmarking various convergence based bGWAS methods.

## PACKAGE DESCRIPTION

We developed hogwash to allow users to perform three bGWAS methods, including an open source implementation of the previously described PhyC algorithm(1), and aggregate genotypes by user-defined groups of mutations. The hogwash function minimally requires a phenotype, a phylogenetic tree, and a set of genotypes. An optional argument may be supplied to facilitate grouping genotypes. The genotypes and tree can be prepared from a multiVCF file by the variant preprocessing tool prewas(17). Hogwash assumes that the genotype is encoded such that 0 refers to wild type and 1 refers to a mutation and that binary phenotypes are encoded such that 0 refers to absence and 1 refers to presence.

In brief, the hogwash workflow (Figure 1A) begins with the user supplying a phenotype, a set of genotypes, and a tree. Hogwash performs ancestral state reconstruction for the phenotype and genotypes to assign phenotype and genotype values to each tree edge (Figure 1B). The interaction of the phenotype with the genotypes is uniquely defined for each of the three association tests. To establish the significance of the interaction the genotypes are permuted and their intersection with the phenotype is recorded as a null distribution. Finally, we introduce an additional metric, *ε*, to capture the interaction between the convergence of the phenotype and genotypes.

### Definitions

To describe the association algorithms, we introduce terms to characterize phenotypes, genotypes, and their interactions. We evaluate node values in a phylogenetic tree through ancestral state reconstruction. *β* is vector where each element corresponds to an edge in this tree.

- 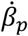 is a binary vector indicating phenotype presence, with a value of 1 for exactly the edges with a child node with value 1 and otherwise 0.
- 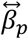 is a binary vector indicating phenotype transitions, with a value of 1 for exactly the edges where the parent differs from the child and otherwise 0.
- 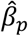 is a continuous vector that has value Δ_*edge*_ = |*phenotype*_*parent node*_ − *phenotype*_*child node*_| for each edge, where Δ_*edge*_ values are normalized from 0 to 1.
- 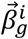 is a binary vector indicating a genotype arising on the tree. It has a value of 1 for exactly the edges where the parent node has value 0 and the child node has value 1, for each genotype *i* in the set of all genotypes.
- 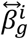 is a binary vector indicating genotype transitions, with a value of 1 for exactly the edges where the parent differs from the child and otherwise 0, for each genotype *i* in the set of all genotypes.
- We define the elementwise sum of *β* as ∑ *β*.

Our three methods use different combinations of *β*_*p*_ and *β*_*g*_. PhyC is concerned with presence and appearance 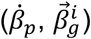. The Synchronous Test is concerned with transitions 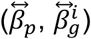. The Continuous Test is concerned with deltas and transitions 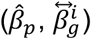.

The interaction of the phenotype and genotypes are summarized as N for each method.

- We define the number of edges where both a genotype arises and the phenotype is present as 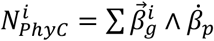, for each genotype *i* in the set of all genotypes.
- We define the number of tree edges where both a genotype changes and the phenotype changes as 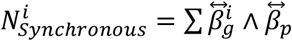, for each genotype *i* in the set of all genotypes.
- We define the sum of the absolute value of phenotype change on only genotype transitions edges as 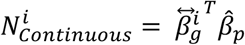, for each genotype *i* in the set of all genotypes.

### PhyC

PhyC is a convergence based bGWAS method introduced by Farhat *et al.*(1) that identified novel antibiotic resistance-conferring mutations in *M. tuberculosis*. To our knowledge, the original PhyC code is not publicly available, but the algorithm is well described in the original paper. The algorithm addresses the following question: Does the genotype transition from wild type, 0, to mutant, 1, occur more often than expected by chance on tree edges where the phenotype is present, 1, than where the phenotype is absent, 0? By requiring the overlap of the phenotype with the genotype transition, instead of genotype presence, associations are not inflated by clonal sampling and thus this approach controls for population structure. We implement the PhyC algorithm as described in Farhat *et al.*(1).

For permutation tests to determine the significance of associations genotype transitions are randomized on the tree with probability proportional to the branch length. The number of edges where the permuted genotype mutation intersects with phenotype presence edges is recorded for each permutation; these permuted 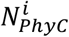 values create a null distribution. An empirical *P*-value is calculated based on the observed 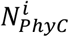 as compared to the null distribution.

Our PhyC implementation (Figure 2) has several important differences from the original paper. First, multiple test correction in hogwash is performed with False Discovery Rate instead of the more stringent Bonferroni correction. Second, hogwash reduces the multiple testing burden by testing only those genotype-phenotype pairs for which convergence is detectable; genotypes with 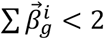 are excluded and genotype-phenotype pairs with 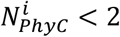 are assigned a *P*-value of 1. Third, ancestral state reconstruction for genotypes and phenotypes was performed using only maximum likelihood. Finally, users sacrifice some robustness in exchange for ease of use by supplying one phylogenetic tree instead of three.

**Figure 2.**
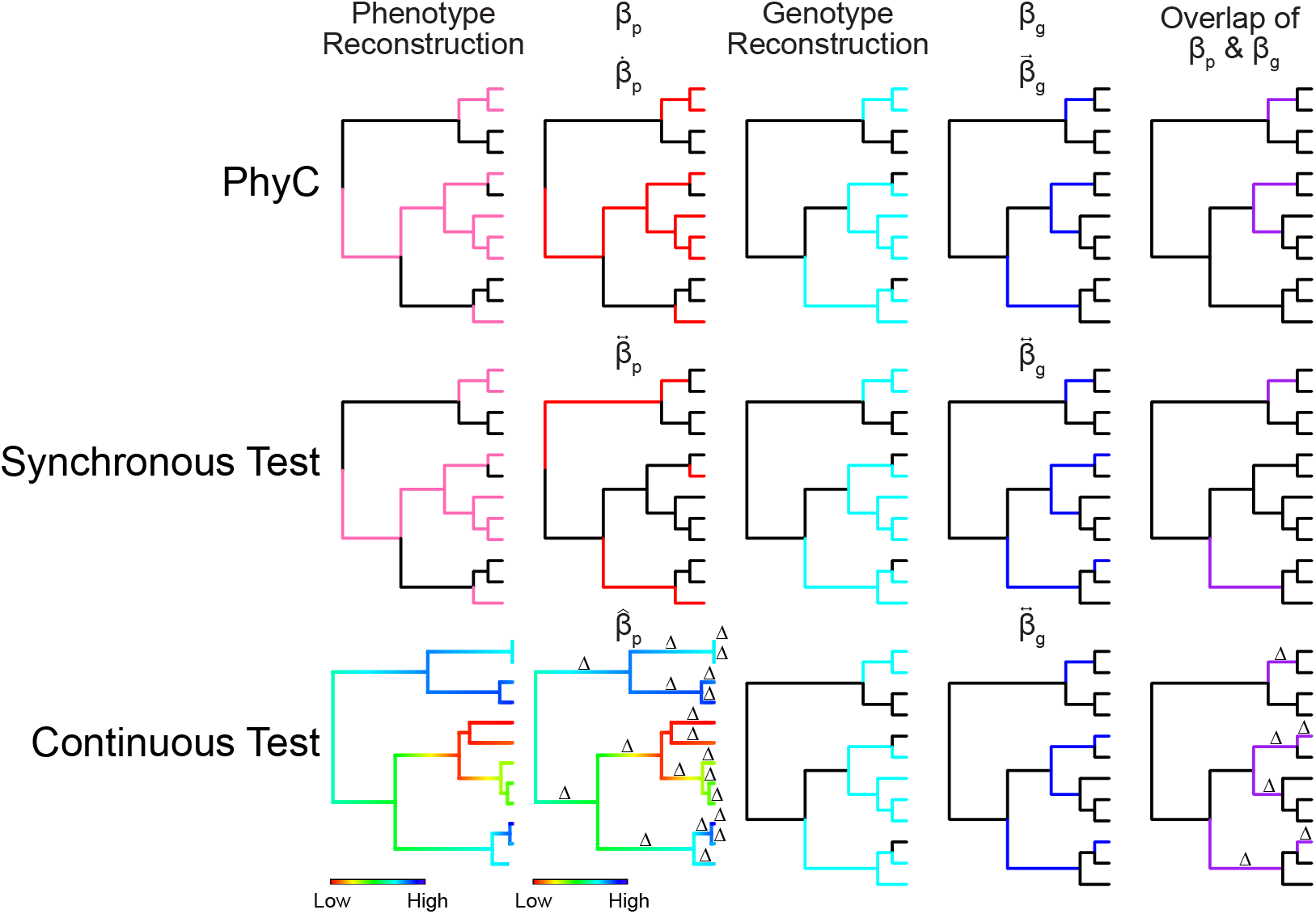
Schematic of PhyC, Synchronous, and Continuous Tests. For all binary trees black indicates 0 and a solid color indicates 1. The Phenotype Reconstruction indicates the ancestral state reconstruction for a simulated phenotype; either binary for PhyC and Synchronous Test or a range of values for the Continuous Test. The *β*_*p*_ indicates the test-specific *β*_*p*_ value taken on each tree edge; 0 or 1 for PhyC and Synchronous Test or the normalized Δ_*edge*_ for the Continuous Test. The Genotype Reconstruction column indicates the ancestral state reconstruction for a simulated genotype; the values are 0 or 1 all algorithms. The *β*_*g*_ indicates the test-specific *β*_*g*_ value taken on each tree edge; the values are 0 or 1 all algorithms. The Overlap of *β*_*p*_ and *β*_*g*_ represents the components of *N*_*test*_ for the specific simulated genotype shown. *β*_*p*_, *β*_*p*_, and *N*_*test*_ are formally described in the Definitions section.

### Synchronous Test

This test (Figure 2) is an extension of PhyC but requires more stringent association between the genotype and phenotype. The Synchronous Test addresses the question: Do genotype transitions occur more often than expected by chance on phenotype transition edges than on phenotype non-transition edges? As in PhyC, the Synchronous Test is only appropriate for binary phenotypes.

Genotypes with 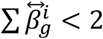 are removed, genotype-phenotype pairs with 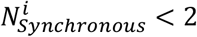 are assigned a *P*-value of 1,and the remaining genotypes are permuted and a null distribution of 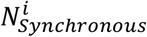 is calculated to determine the significance of each genotype.

This test is similar to the Simultaneous Score in treeWAS(9). The Simultaneous Score is derived from the number of edges on the tree where the genotype and phenotype transition in the same direction (both have a parent node of 0 and an inferred child node of 1 or both have a parent node of 1 and child node of 0). In contrast, our newly developed Synchronous Test allows for the phenotype and genotype transition directions to mismatch, thus allowing for a genotype to have opposing effects on a phenotype. Such opposing effects of a genotype on a phenotype could arise when grouping mutations in the same gene that differentially impact gene function, or even for an individual mutation whose phenotypic impact may be dependent on genetic background.

### Continuous Test

The Continuous Test (Figure 2) is a novel application of a convergence based GWAS method to continuous phenotypes. The Continuous Test addresses the question: Does the phenotype change more than expected by chance on genotype transition edges than on genotype non-transition edges?

As above, the genotypes with 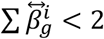 are removed; the remaining genotypes are permuted and a null distribution of the 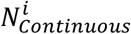 is calculated to determine the significance of each genotype.

### User inputs

The user must provide a phylogenetic tree, a set of genotypes, and a phenotype. The user may optionally provide a key that maps individual genomic loci into groups in order to use hogwash’s grouping feature. For a detailed description of the user inputs please see the Supplementary Package Description.

### Hogwash outputs

The package produces two files per test: data (.rda) and plots (.pdf). The data file contains many pieces of information, including *P*-values for each tested genotype. The plots are described below in the results section.

### Grouping feature

To identify an association between a genomic variant and a phenotype hogwash requires that a variant occur in multiple different lineages. Hogwash may classify some causal variants as independent of a phenotype if they are weakly penetrant. To surmount this issue, related genomic variants may be aggregated to capture larger trends at the grouped level. For example, a user may apply this method to group only nonsynonymous SNPs by gene to use hogwash to detect associations between the mutated gene and the phenotype. Grouping related variants can improve power through a reduction in the multiple testing correction penalty. However, the power benefits are dependent on grouping variants with similar effect directions.

By default, hogwash implements the grouping features by first performing ancestral state reconstruction for each individual locus (Figure 3). Then those loci are joined as indicated in the user supplied key. Grouped loci with 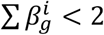 are excluded from analysis. After this point hogwash runs as previously described for non-grouped genotypes. Alternatively, users may group together related genomic variants prior to ancestral reconstruction (Supplementary Methods). The two grouping approaches are compared in Figure S1.

**Figure 3.**
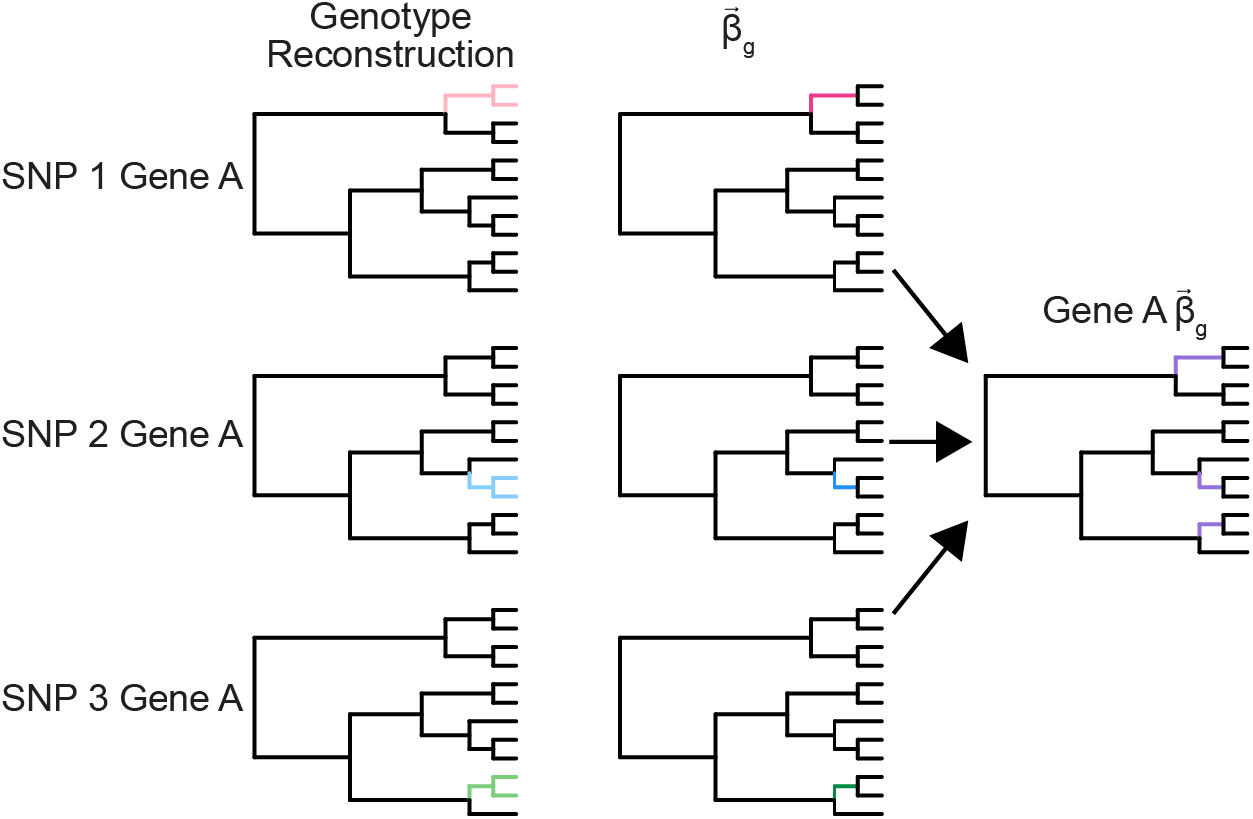
Example of hogwash grouping feature on simulated data. In this case, three SNPs are found in the same gene (Gene A). No individual SNP is convergent on the tree. Hogwash performs ancestral state reconstruction on each SNP. The edges where SNP presence is inferred are colored. Next, hogwash identifies the 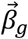 for each SNP (colored edges). Finally, hogwash combines the three SNP 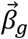 together to create the Gene A 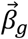 (purple edges). When the SNPS are grouped into Gene A the genotype converges on the tree. In this example, pre-ancestral reconstruction and post-ancestral reconstruction grouping results are identical. See Figure S1 for scenarios illustrating differences in the two grouping approaches.

## METHODS

### Data simulation

#### Trees

We simulated eight random coalescent phylogenetic trees with 100 tips each; four trees were used for the Continuous Test and four trees were used for the binary tests.

##### Tree edge filtering

Low confidence edges are defined as those edges with low bootstrap support (default <70%), those that are more than 10% of the total tree length, or those with low genotype or phenotype ancestral reconstruction support (maximum likelihood <0.875). Low confidence edges are ignored during permutation testing.

#### Phenotypes

##### Motivation for simulating phenotypes under two evolutionary models

For each tree we simulated phenotypes under different evolutionary models: either Brownian motion or white noise. A phenotype modeled well by Brownian motion follows a random walk along the tree. A phenotype modeled well with white noise appears to be independent of tree structure and may suggest a role for horizontal gene transfer, gene loss, or convergent evolution(18). A white noise phenotype may be better suited to the hogwash algorithms than a phenotype modeled by Brownian motion given the requirement for phylogenetic convergence.

##### Calculation of phylogenetic signal

Phylogenetic signal is a metric that captures the tendency for closely related samples on a tree to be more similar than random samples. Phylogenetic signal is calculated by different metrics for continuous and binary traits; continuous traits are measured by *λ* while binary traits are measured by *D* (Figure S2). A continuous phenotype that is modeled well by Brownian motion has a phylogenetic signal, *λ*, near 1 while a white noise phenotype has a phylogenetic signal near 0(19). In contrast, a binary phenotype that is modeled well by Brownian motion has a phylogenetic signal, *D*, near 0 while a white noise phenotype has a phylogenetic signal near 1(20).

##### Simulation of phenotypes on trees

For each tree we simulated four phenotypes fitting a Brownian motion model and four phenotypes fitting a white noise model. For phenotypes modeling Brownian motion, binary phenotypes were restricted to −0.05 < *D* < 0.05 and continuous phenotypes to 0.95 < *λ* < 1.05. For phenotypes modeling white noise, binary phenotypes were restricted to 0.95 < *D* < 1.05 and continuous phenotypes to −0.05 < *λ* < 0.05.

#### Genotypes

For each simulated tree a set of unique binary genotypes were generated. We generated genotypes that span a range of phylogenetic signals, degree of similarity to the phenotype, and prevalence.

##### Genotypes to be used in PhyC and the Synchronous Test

First, 25,000 binary genotypes were generated using ape::rTraitDisc; these genotypes have a range of phylogenetic signals(21). Second, these genotypes were duplicated and randomized with the following approach: one quarter had 10% of tips changed, one quarter had 25% of tips changed, one quarter had 40% of tips changed, and one quarter were entirely redistributed. Third, we removed any simulated genotypes present in 0, 1, *N* − 1, or *N* samples. Fourth, we subset the genotypes to keep only unique presence/absence patterns. Fifth, we subset genotypes to only those within a range of −1.5 < *D* < 1.5. These filtering steps result in a reduction in the data set size (range 2214-2334).

##### Genotypes to be in used in the Continuous Test

In addition to the five steps above we added two more data generation steps. First, we made all possible genotypes based on the rank of the continuous phenotype. Second, we made genotypes based on which edges of the tree had high Δ_*edge*_. The filtering steps reduced the data set size (range 1234-1310).

### Hogwash on simulated data

We ran hogwash for each of the tree-phenotype-genotype sets. In addition to generating *P*-values for each tested genotype, hogwash also reports convergence information. We ran hogwash with the following settings for single locus analysis: permutations = 50,000; false discovery rate = 0.0005 (binary), 0.05 (continuous); bootstrap value = 0.70; no genotype grouping key was provided. For grouped analyses the settings were identical except that a grouping key was generated, and hogwash was run with both grouping methods (pre- and post-ancestral reconstruction). For the grouped analyses only PhyC was run on simulated Brownian motion phenotype 1, simulated genotype 1, and simulated tree 1. The grouping key assigned each approximately 3 unique simulated variants to each created “gene;” resulting in approximately one-third as many input genotypes when compared to the single locus analysis.

#### Calculation of *ε*

We introduce *ε* to quantify the degree of shared phenotype convergence and genotype convergence. Low values of *ε* indicate a lack of overlap in the edges where the phenotype and genotype converge. High values of *ε* indicate many instances of overlap in the edges were the phenotype and genotype converge. By reducing these patterns of convergence into a simple number, *ε*, we can more easily contextual convergence based bGWAS results. We define an *ε* for each algorithm.

- 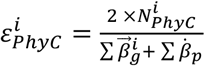, for each genotype *i* in the set of all genotypes.
- 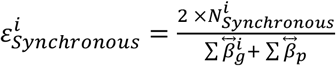, for each genotype *i* in the set of all genotypes.
- 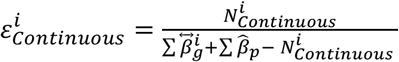, for each genotype *i* in the set of all genotypes.
- For each *ε*, 0 ≤ *ε* ≤ 1.

### Data analysis

Statistical analyses were conducted in R v3.6.2(22). The R packages used can be found in the simulate_data.yaml file on GitHub(21,23–27) and can be installed using miniconda(28).

## RESULTS

### Motivation for evaluating Hogwash on simulated data

Given our lack of comprehensive knowledge of the genetic variation contributing to any phenotype, it is not feasible to quantify sensitivity/specificity on real data. We therefore generated data that simulates genotype and phenotype distributions covering a spectrum of realistic evolutionary scenarios (spanning Brownian motion to white noise). Our goal is not to validate the premise of using convergence based approaches, as these have been previously shown to provide useful biological insights, but rather to understand how our approach detects convergence for phenotypes with different evolutionary regimes. The following analyses can guide users in the appropriate use cases and the applicability of this method to their data. Particularly, these results provide context for interpreting the strength of the observed associations.

### Hogwash output for simulated data

Hogwash outputs two sets of results: a data file and a PDF file with plots. Each run of PhyC produces at least three plots: the phenotype reconstruction (Figure 4A), a Manhattan plot (Figure 4D), and a heatmap of genotypes (Figure 4E). The phenotype reconstruction is highlighted on the tree (Figure 4A). The Manhattan plot shows the distribution of *P*-values from the hogwash run (Figure 4D). The heatmap shows the genotype reconstruction and phenotype reconstruction for each tree edge (rows) and genotype (columns) (Figure 4E). The genotypes are clustered by the presence/absence pattern. Two additional plots are produced for each genotype that is significantly associated with the phenotype: a phylogenetic tree showing the genotype transition edges (Figure 4B) and the null distribution of N_*phyc*_ (Figure 4C). The two grouping approaches identified novel associations missed when PhyC was run without a grouping key (Figure S3).

**Figure 4.**
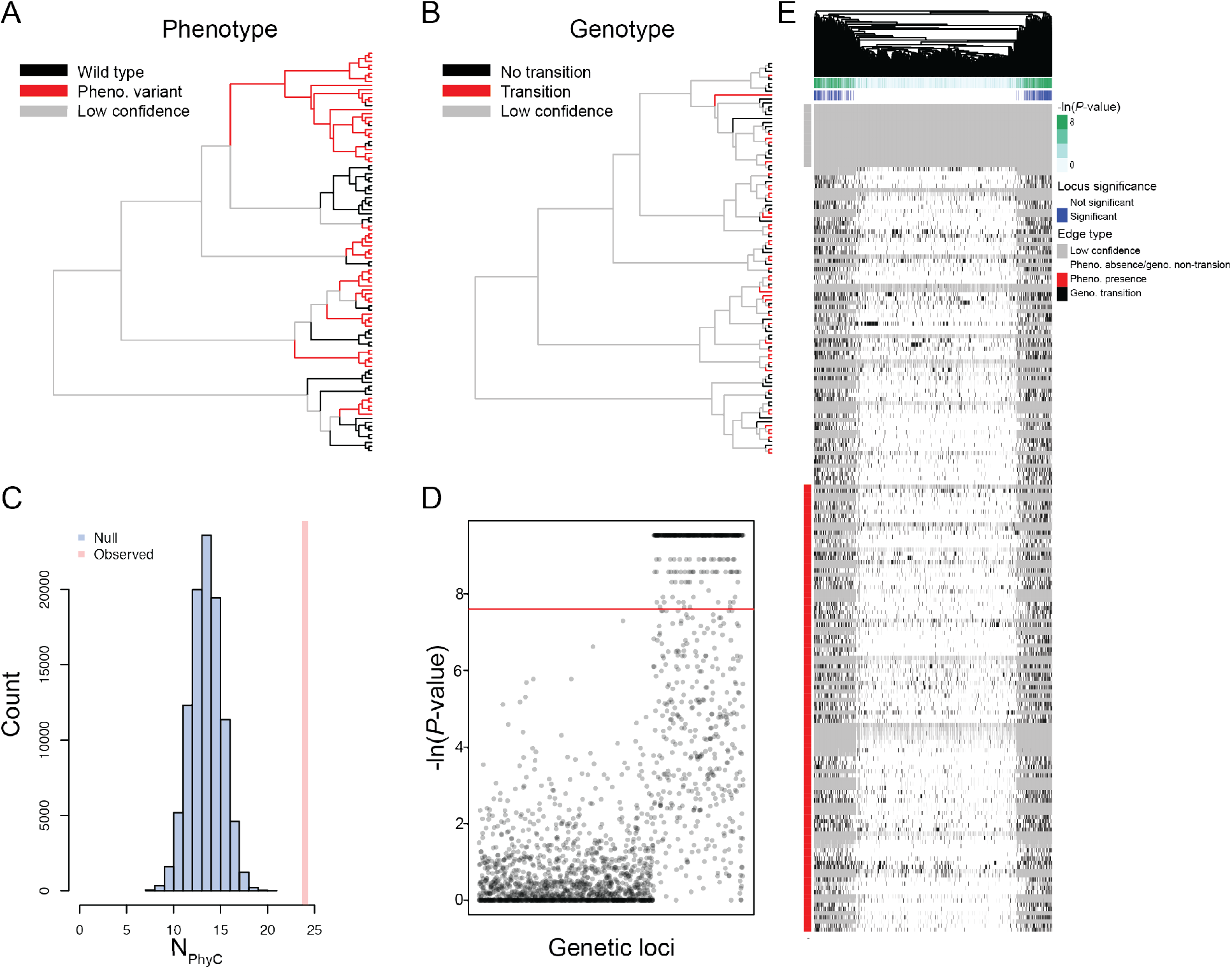
Example output from hogwash PhyC results from simulated data. A) Phenotype reconstruction. Edges with: phenotype presence in red; phenotype absent in black; low confidence in tree or low confidence phenotype ancestral state reconstruction in gray. B) Genotype transitions. Edges with: genotype mutations that arose in red; genotype mutation did not arise in black; low confidence in tree or low confidence genotype ancestral state reconstruction in gray. C) Null distribution of N_*phyC*_. D) Manhattan plot. The genetic loci were simulated to achieve a range of phylogenetic signals. The left most two-thirds of genetic loci were simulated under Brownian motion models (mean *D*=0.16) while the remain third were modeled by white noise (mean *D*=0.99). E) Heatmap with tree edges in the rows and genotypes in the columns. The genotypes are hierarchically clustered. The genotypes are classified as being a transition edge in black or non-transition edge in white. The column annotations pertain to loci significance; green indicates the *P*-value while blue indicates that the *P*-value is more significant than the user-defined threshold. The row annotation classifies the phenotype at each edge; red indicates phenotype presence and white indicates phenotype absence. Gray indicates a low confidence tree edge; low confidence can be due to low phenotype ancestral state reconstruction likelihood, low genotype ancestral state reconstruction likelihood, low tree bootstrap value, or long edge length.

The Synchronous Test and Continuous Test output plots that reflect their test-specific *β* and N definitions (Figure S4, S5). Running hogwash on 100 samples required <3 hours and <2 GB of memory for binary data and <5 hours and <2 GB of memory for continuous data (Figure S6).

### Hogwash evaluation on simulated data

To help users identify optimal use cases and also interpret hogwash results we describe the behavior of hogwash on simulated data. We note that this assessment is not meant to convey performance in the sense of calculating sensitivity and specificity, but rather evaluate whether hogwash can robustly detect the association between phenotypic and genotypic convergence. To guide our assessment, we compared the relationship between the *P*-value and *ε* values produced by hogwash on sets of simulated data constructed using different evolutionary models (Figure 5). *ε* is a quantification of the relationship between phenotype convergence and genotype convergence. Low *ε* values indicate little to no intersection of phenotype convergence and genotype convergence, while higher *ε* values indicate their increased intersection. The *ε* value is always a fraction between 0 and 1 and therefore obscures information about the sample size; to account for the number of samples in the tree we recommend always interpreting *ε* value for any locus with its *P*-value.

**Figure 5.**
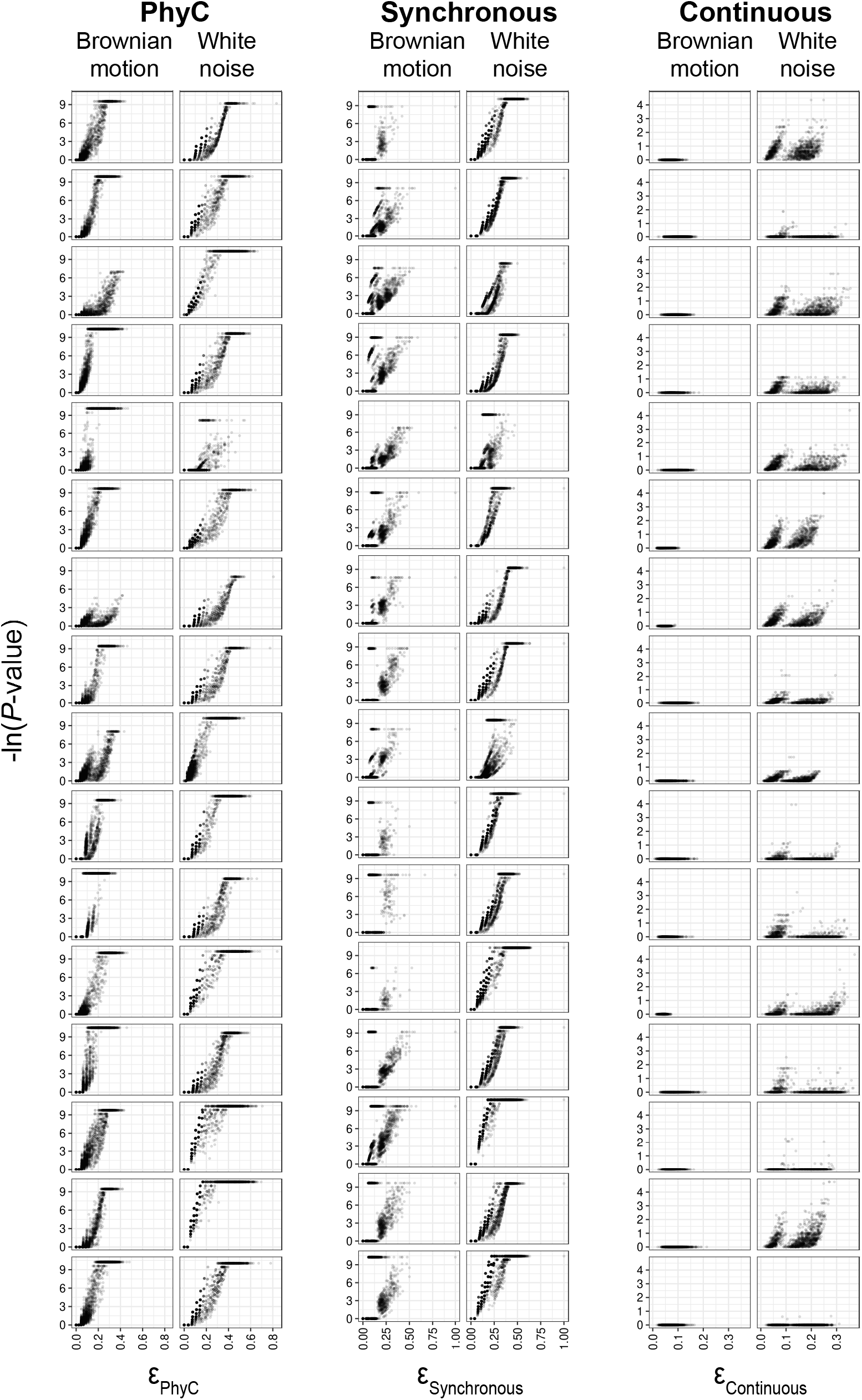
High *ε* values correlate with increased significance. Each plot is a tree-phenotype pair. Each point represents one genotype-phenotype pair. Brownian motion and White noise refer to the evolutionary regime modeled by the phenotype. The genotypes span a range of phylogenetic signals.

For binary phenotypes, we observe an overall strong positive association between −log(*P*-value) and *ε*, demonstrating that as the intersection of phenotype convergence and genotype convergence increase hogwash predicts that it is less likely that they intersect due to chance (Table 1). In other words, below a certain *ε*_*binary*_ threshold (*ε*_*binary*_ is *ε*_*phyC*_ or *ε*_*Synchronous*_), hogwash attributes the association between the genotype convergence and phenotype convergence to chance; from Figure 5 the user can get a sense for the range of this *ε*_*binary*_ threshold under different evolutionary regimes.

**Table 1.**
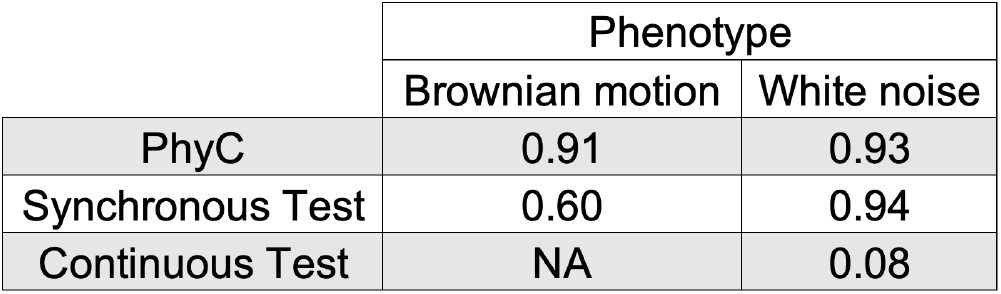
Mean Spearman’s rank correlation coefficient for −ln(*P*-value) versus *ε* from hogwash run on simulated data. The ρ could not be calculated for the results from the Continuous Test on the Brownian motion phenotypes because, after multiple testing correction, all *P*-values are identical.

|  | Phenotype |  |
| --- | --- | --- |
|  | Brownian motion | White noise |
| PhyC | 0.91 | 0.93 |
| Synchronous Test | 0.60 | 0.94 |
| Continuous Test | NA | 0.08 |

For the simulated continuous data an *ε*_*Continuous*_ threshold that separates meaningful genotype-phenotype associations from associations by chance is less apparent. Higher *ε*, low significance values demonstrate that some overlap of *β*_*g*_ and *β*_*p*_ is likely by chance given the data. Low *ε*, high significance genotype-phenotype pairs demonstrate that sometimes small amounts of *β*_*g*_ and *β*_*p*_ overlap are unlikely, however that does not necessarily suggest that these hits are the best candidates for *in vitro* follow up. We suspect that these associations are largely driven by poor exploration of the sampling space, despite running many permutations, because of the edge-length based sampling probability of the permutation method. Therefore, it is essential *P*-values be interpreted within the context of *ε*. Notably, the Continuous Test was only able to detect significant genotype-phenotype associations for phenotypes modeled by white noise, suggesting this method is particularly sensitive to the phenotype’s evolutionary model. We observe for both the binary and continuous tests that *ε* is more tightly correlated with −log(*P*-value) for phenotypes characterized by white noise than by Brownian motion (Table 1), indicating that hogwash performs better under a white noise model. Therefore, we suggest using the report_phylogenetic_signal function on the phenotype prior to running hogwash to ascertain the appropriateness of these algorithms for the dataset.

## DISCUSSION

We have developed two algorithms for convergence based bGWAS that are particularly well suited for phenotypes modeled by white noise. Hogwash, is straightforward to install in R, accepts easy-to-format data inputs (described in detail on the wiki), and provides publication ready plots of the GWAS results. Hogwash also introduces grouping features to aggregate related genomic variants to increase detection of convergence for weakly penetrant genotypes. Hogwash is best used for datasets comprising binary and/or continuous phenotypes, phenotypes fitting white noise models, situations where convergence may occur at the level of genes or pathways and with datasets whose size can be accommodated given the time and memory constraints of convergence methods.

The results of running hogwash on simulated data suggest that after a certain *ε* threshold, it unlikely that the intersection between phenotype convergence and genotype convergence occurs by chance, particularly for white noise phenotypes. Given the variability in results within each method, as shown in Figure 5, users may want to contextualize the statistical significance of the tested genetic loci with the amount of convergence possible for any one particular data set; to facilitate this the hogwash output includes both *P*-values and *ε*.

The simulated data set presented here is published to serve as a resource or template for future work focused on benchmarking convergence based bGWAS software as such a dataset has not yet, as far as we are aware, been made available(29). The simulated data set is available on GitHub and includes convergence information for each phenotype, genotype, and their intersection.

## AUTHOR CONTRIBUTIONS

KS and ESS conceptualized the project and edited the manuscript. KS designed and implemented the software, performed the analysis, prepared the original draft, and visualized the data. ESS supervised the project.

## ACKNOWLEDGEMENTS

We thank Brad Saund for his help formalizing the continuous algorithm *ε* definition.

