## Supplement for "hogwash: Three Methods for Genome-Wide Association Studies in Bacteria"

### Extended Package Description

#### User Provided Inputs

*Tree*

The provided phylogenetic tree may be rooted; unrooted trees will be midpoint rooted. The tree must be fully bifurcating. We recommend building the phylogenetic tree with an outgroup using a maximum likelihood framework, root to the outgroup, and then remove the outgroup prior to running hogwash. We also recommend that trees be built from recombination-filtered genomic data(1).

*Genotype*

The required structure of the genotype data object is a matrix. The rows correspond to samples and should be ordered to match the tips of the phylogenetic tree. The row names must exactly match the tree's tip labels. The columns correspond to individual genotypes. The matrix should have both row names and column names. Genotypes can be SNPs (core genome), genes (accessory genome) or other types (indels, pathways, etc...). Genotypes must be encoded in binary (0/1). We assume that the user has chosen a meaningful reference and recoded the reference allele as 0 and variants as 1.

*Phenotype*

The required structure of the phenotype data object is a matrix. The rows correspond to samples and should be ordered to match the tips of the phylogenetic tree. There should only be one column, which contains the phenotype data. The matrix should have both row names and a column name. The row names must exactly match the tree's tip labels. The phenotype can either be binary (0/1) or continuous. We assume that the user has chosen a meaningful reference for binary phenotypes.

*Optional: Genotype Grouping Key*

To perform grouping, for example of SNPs into genes, the user must provide a key that maps the individual loci into groups. This matrix has two columns, where the first column has names of the genotypes included in the provided genotype matrix. The second column has names for the group to which the item in the first column belongs. If the individual locus should be mapped to more than one group, the user should add additional rows for each additional group. Row names are not required. The column names are used in output plots and therefore must be included.

#### Filters and checks performed on user-provided inputs

*Genotype*

Hogwash filters will remove genotypes whose variants are present in only $0$, $1$, $N$ or $N-1$ genomes because convergence is not detectable. This removal step occurs prior to ancestral state reconstruction for the non-grouped test or after grouping in the grouped test. Additionally, variants with missing information are removed as are variants with fewer than two high confidence transition edges because otherwise no convergence is detectable ($\beta_g < 2$). Any edges with low genotype ancestral state reconstruction support (maximum likelihood < 0.875) are excluded from the analysis.

*Phenotype*

Hogwash filters will exclude any edges with low phenotype ancestral state reconstruction support (maximum likelihood < 0.875) from the analysis. Hogwash reports the phenotype's phylogenetic signal to the user. The user can check the phylogenetic signal prior to running hogwash with the function report_phylogenetic_signal.

*Tree*

Hogwash hard filters exclude the following tree edges from analysis: A) low bootstrap support (default: <70%) and B) overly long (>10% of total tree edge length). Overly long edges are excluded because ancestral state reconstruction accuracy decreases with edge length(2). The tree must be fully bifurcating. Unrooted trees are midpoint rooted.

*Grouping*
The user can choose between two grouping methods: either pre- or post-ancestral reconstruction grouping (Figure S1). Pre-ancestral reconstruction grouping occurs before the genotype ancestral states are inferred; genotypes are grouped, and then ancestral reconstruction is performed. Pre-ancestral reconstruction grouping is fast but is not as sensitive as post- ancestral reconstruction grouping. Post-ancestral reconstruction grouping occurs after the ancestral states and genotype transitions are determined for each individual (un-grouped) genotype.

### Supplementary Figures

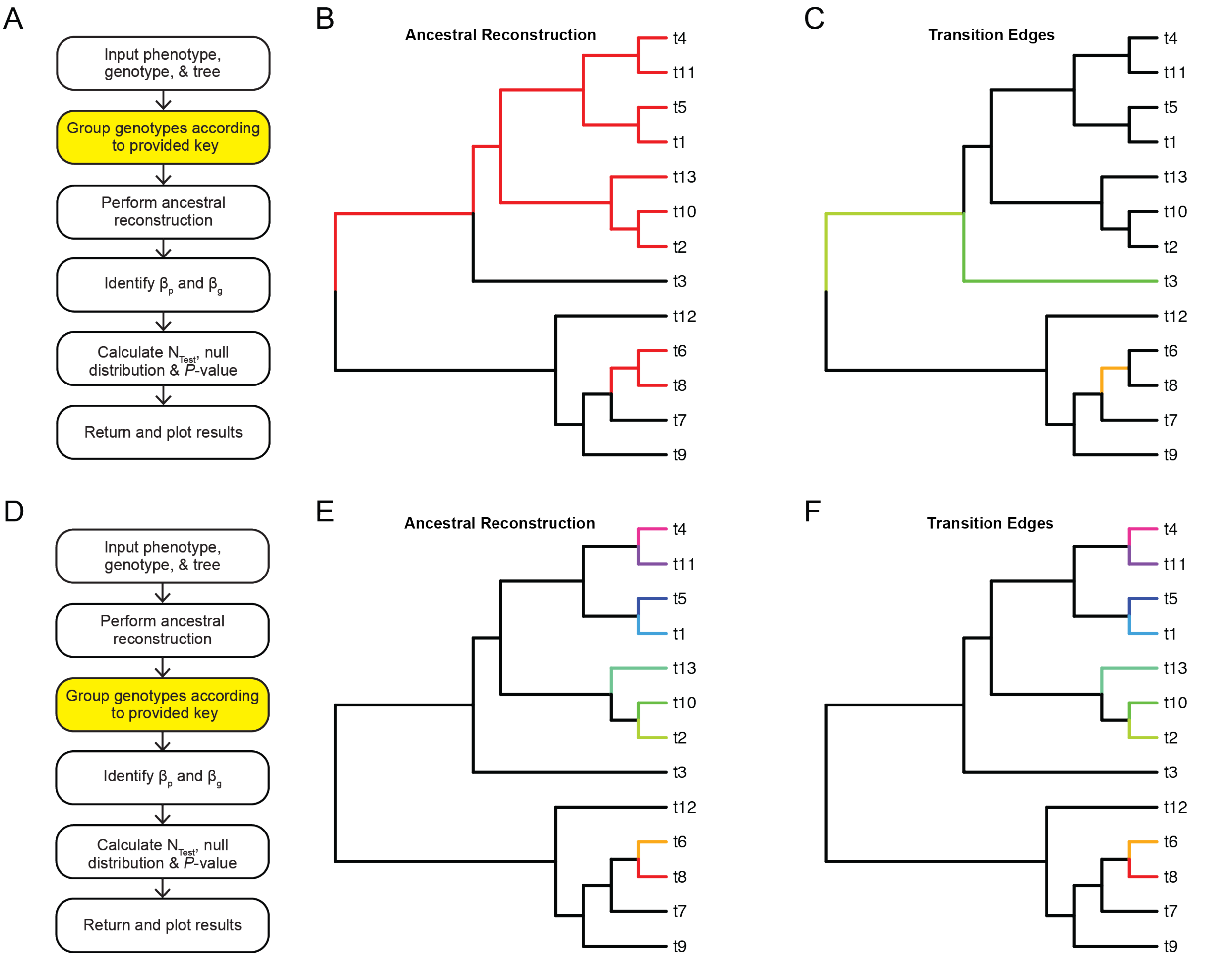


**Figure S1. Pre-ancestral reconstruction grouping versus post-ancestral reconstruction grouping.** On a simulated tree 9 unique SNPs were assigned to tips t4, t11, t5, t1, t13, t10, t2, t6, and t8. In this scenario all 9 SNPs are found in Gene A. A) Hogwash workflow for the pre-ancestral reconstruction grouping method. B) The ancestral state reconstruction results for the Gene A SNPs resulting from pre-ancestral reconstruction grouping. C) Given the ancestral reconstruction results in B, there are 3 genotype transition edges. D) Hogwash workflow for the post-ancestral reconstruction grouping method. E) The ancestral state reconstruction results for the 9 SNPs found in Gene A. F) Given the ancestral reconstruction results in D, there are 9 genotype transition edges.

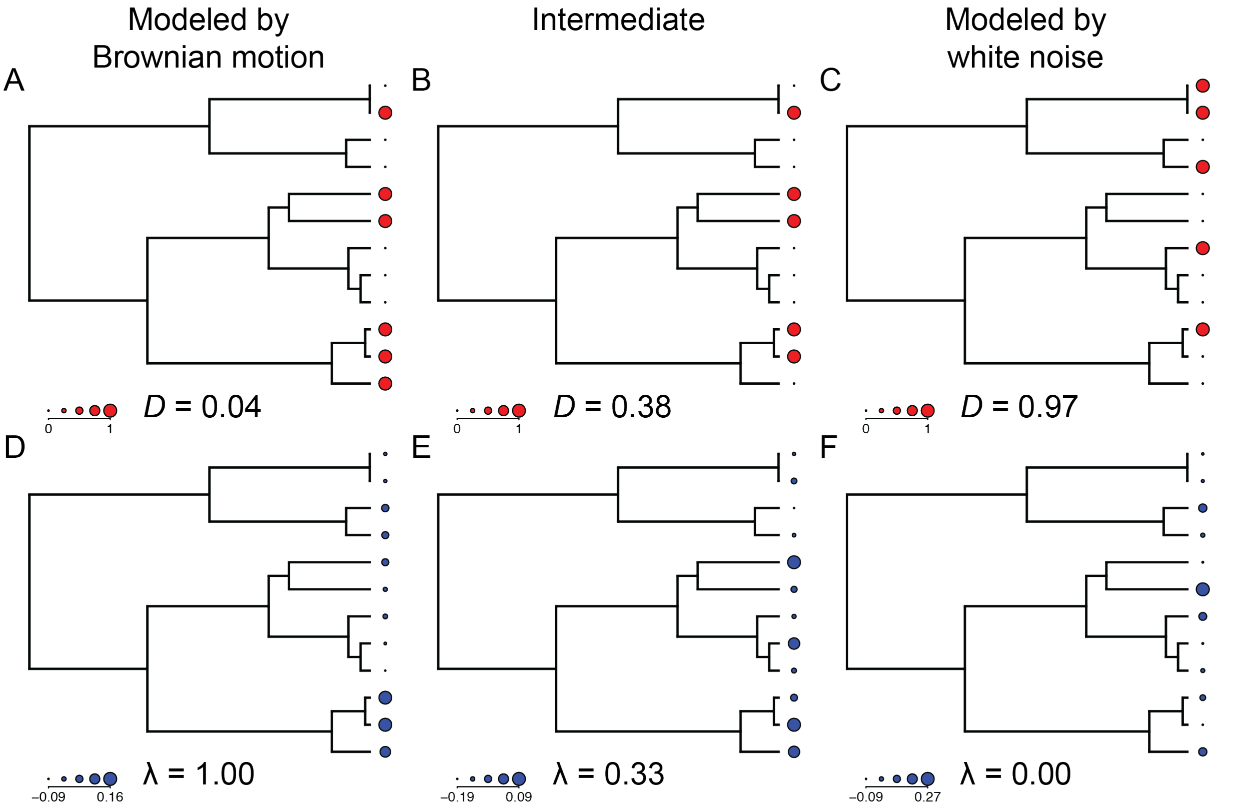


**Figure S2. Phylogenetic signals calculated on simulated trees and phenotypes.** A-C) The phylogenetic signal for the binary phenotypes in blue calculated as *D*. D-F) The phylogenetic signal for the continuous phenotypes in blue calculated as *λ*. Data in A & D fit a Brownian motion model. Data in B & E have phylogenetic signals intermediate between a Brownian motion model and white noise model. Data in C & F fit a white noise model.

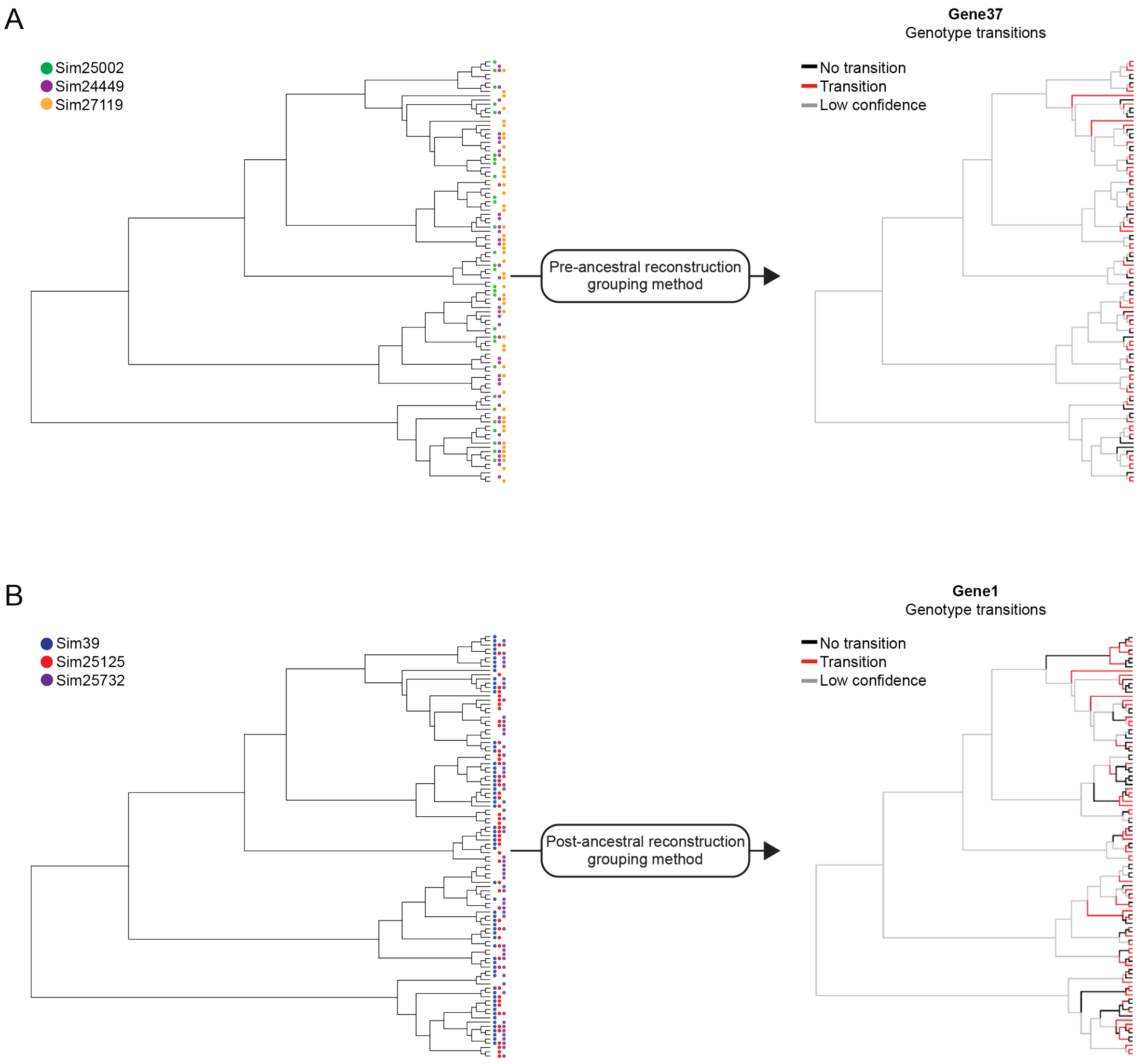


**Figure S3. Selected hogwash grouped PhyC results using both grouping methods.** A) Results from the pre-ancestral reconstruction grouping method. Three SNPs were selected to combine into Gene37. The left tree shows the presence of each SNP. When run through PhyC as individual loci the SNPs had the following -log(*P*-values): Sim25002 = 3.8, Sim2449 = 3.7, Sim27119 = 3.6. The right tree shows the Gene37 transitions. Gene37 had -log(*P*-value) = 7.1. B) Results from the post-ancestral reconstruction grouping method. Three SNPs were selected to combine in Gene1. The left tree shows the presence of each SNP. When run through PhyC as individual loci the SNPs had the following -log(*P*-values): Sim39 = 3.5, Sim25125 = 3.2, Sim25732 = 3.2. The right tree shows the Gene1 transitions. Gene1 had -log(*P*-value) = 9.3.

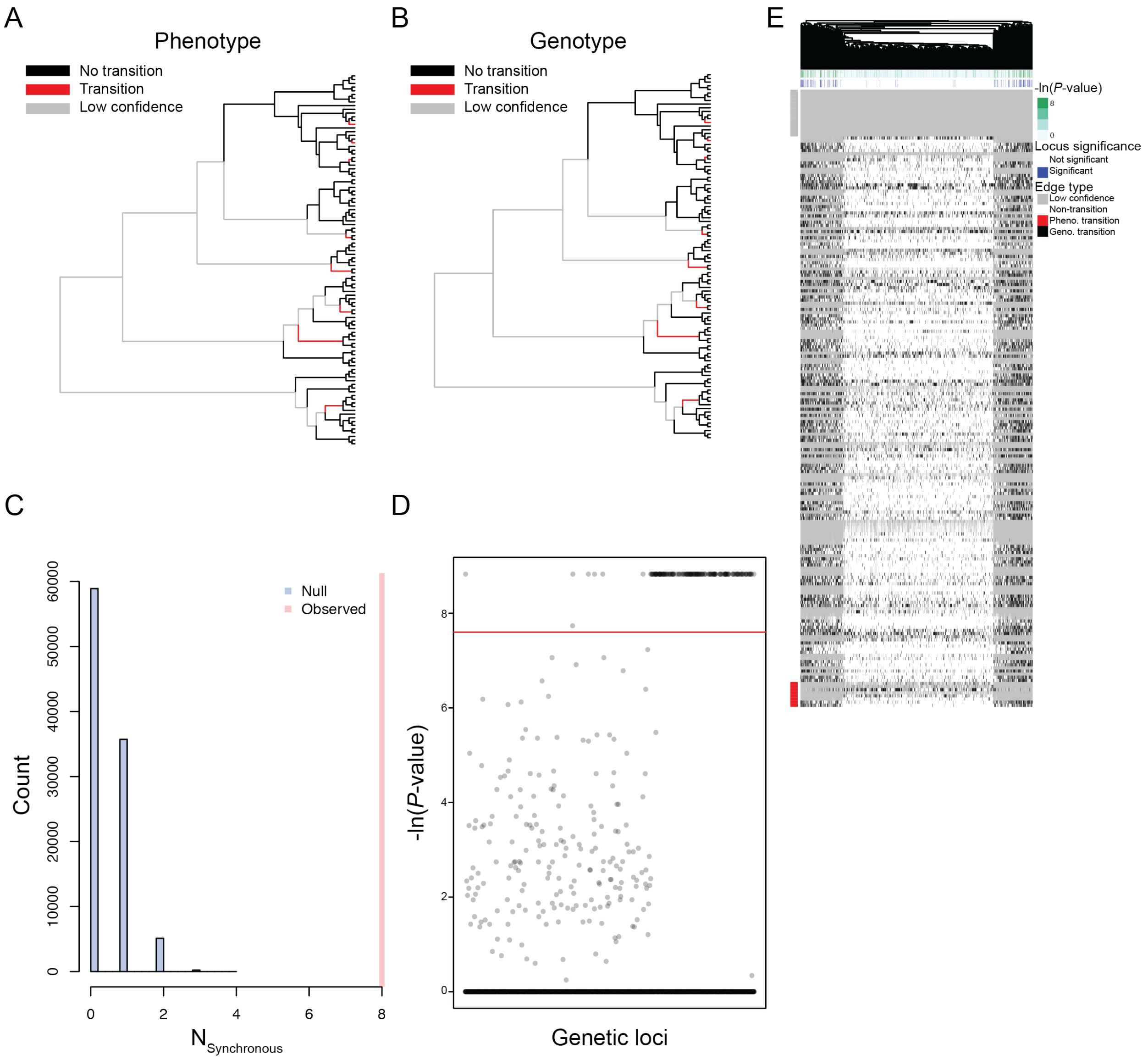


**Figure S4. Example output from hogwash Synchronous Test results on simulated data.** A) Phenotype transitions. Edges with: phenotype transition in red; no phenotype transitions in black; low confidence in gray. B) Genotype transitions. Edges with: genotype transitions in red; non-transition in black; low confidence in gray. C) Null distribution of $N_{Synchronous}$; observed value in red. D) Manhattan plot. Significance threshold indicated in red. Genotypes have a range of phylogenetic signals. E) Heatmap with tree edges in the rows and genotypes in the columns. The genotypes are hierarchically clustered. The genotypes are classified as being a transition edge in black or non-transition edge in white. The column annotations pertain to the *P*-value; green is the *P*-value and blue indicates that the *P*-value is more significant that the user-defined threshold. The row annotation classifies the phenotype edge type. Phenotype transition edges are in red and non-transition edges are in white. Gray indicates a low confidence tree edge; low confidence can be due to low genotype ancestral state reconstruction likelihood, low tree bootstrap value, or long edge length.

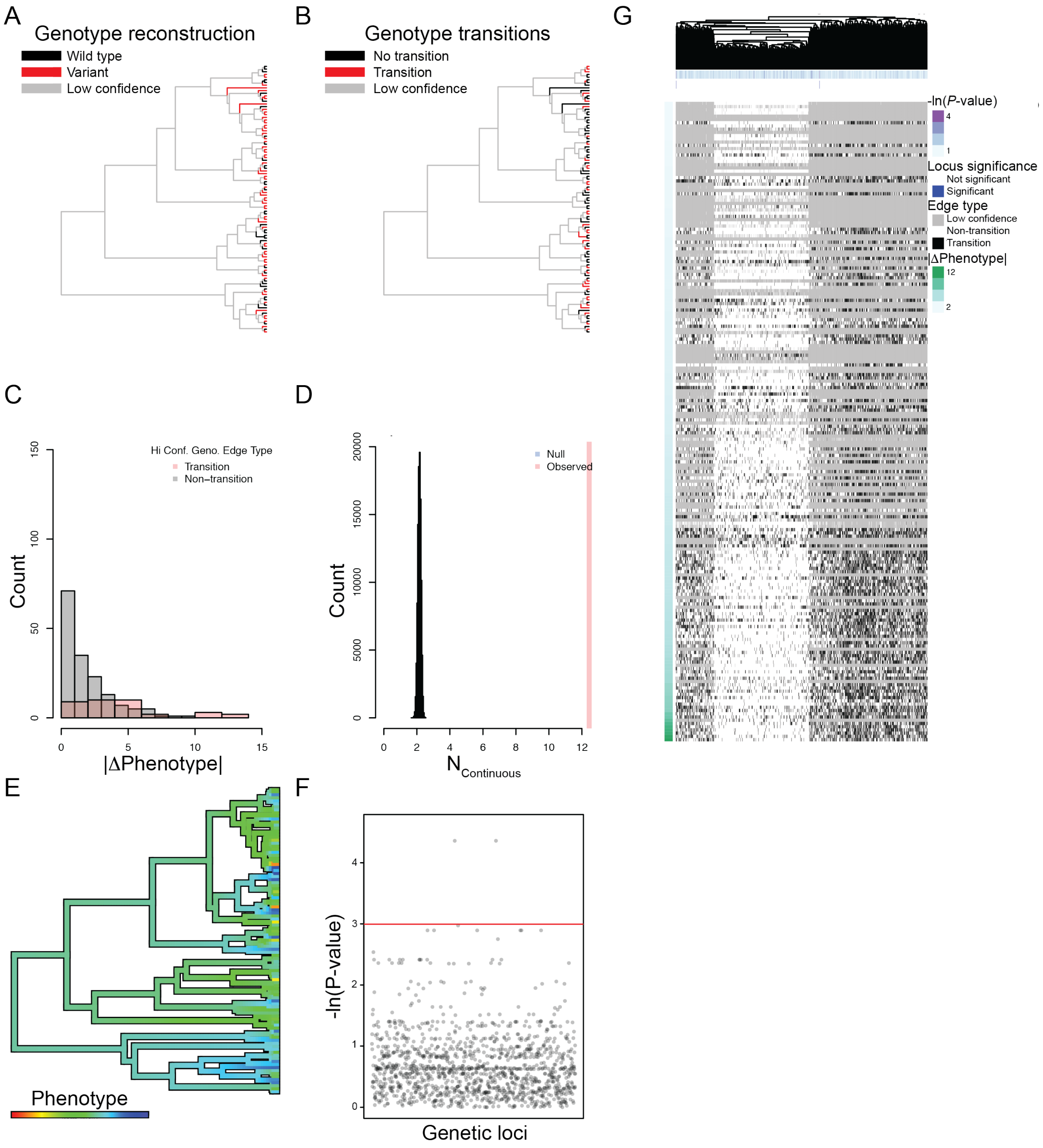


**Figure S5. Example output from the Continuous Test run on simulated data.** A) Reconstruction for a simulated genotype. Wild type in black, variant presence in red, and low confidence edges in gray. B) Genotype transition edges in red; non-transition edges in black; low confidence in gray. C) Histogram of the change in phenotype per edge for high confidence tree edges. Genotype transition edges in red and genotype non-transition edges in gray. D) Null distribution of $N_{Continuous}$; observed value in red. E) Ancestral reconstruction of phenotype. F) Manhattan plot. The significance threshold is indicated in red. Genotypes have a range of phylogenetic signals. G) Heatmap with tree edges in the rows and genotypes in the columns. The genotypes are hierarchically clustered. The genotypes are classified as being a transition edge in black or non-transition edge in white. The column annotations pertain to the *P*-value; purple indicates the *P*-value and blue indicates that the *P*-value is more significant that the user-defined threshold. The row annotation shows the absolute value in the phenotype change per edge. Gray indicates a low confidence tree edge; low confidence can be due to low genotype ancestral state reconstruction likelihood, low tree bootstrap value, or long edge length.

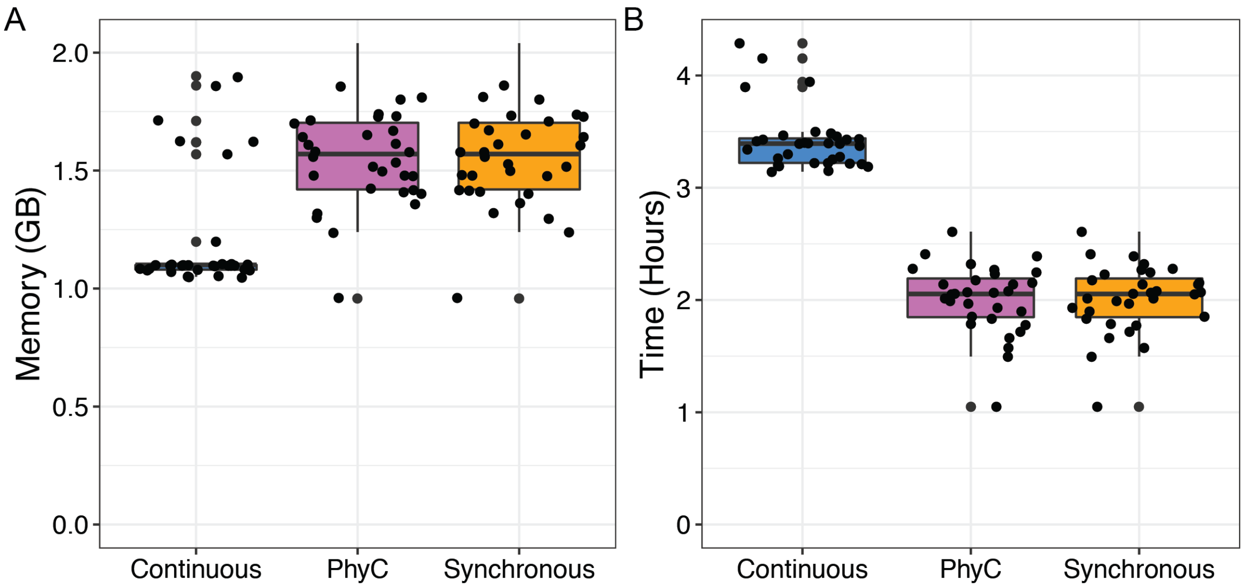


**Figure S6. Memory usage and run time for hogwash on simulated data.**
